# Odonata diversity of the eastern region of Bangladesh with four new addition to the Bangladeshi dragonfly fauna

**DOI:** 10.1101/195438

**Authors:** Md Kawsar Khan

**Affiliations:** Department of Biochemistry and Molecular Biology, School of Life Sciences, Shahjalal University of Science and Technology, Sylhet-3114, Bangladesh; Department of Biological Sciences, Macquarie University, Sydney, New South Wales-2109, Australia

**Keywords:** Bangladesh, Dragonfly, Damselfly, Indo-Burma biodiversity hotspot, Odonata diversity

## Abstract

A study was conducted in the eastern region to contribute to the Odonata fauna of Bangladesh. A total of 76 species belonging to 9 families have been recorded during the study period of April 2014 to July 2016. Three Zygopteran species e.g, *Ceriagrion rubiae*, *Calicnemia imitans*, *Prodasineura autumnalis* and one Anisopteran species e.g, *Megalogomphus smithii* have been newly discovered from Bangladesh. *Megalogomphus* genus have been first time recorded from Bangladesh.

## 1.0 Introduction

Bangladesh, situated in the South East Asia, possesses an enormous area of wetlands including ponds, rivers, freshwater lakes and marshes along with extensive mangrove swamps. Moreover, the hilly areas of the north east and south east region receive precipitation throughout the year and are rich in waterfalls and streams. In addition, during monsoon many paddy fields and irrigation channels hold water more than three months and generate numerous temporary water reservoirs. This diverse range of water resources offers ambient microhabitat for many Odonata species (Chowdhury & Mohiuddin 1994). Till date, 103 species of Odonates have been recorded from Bangladesh (Begum et al. 1977; Chowdhury & Akhteruzzaman 1983; Chowdhury & Mia 1989; Chowdhury & Mohiuddin 1993; Noruma & Alam 1995; Chowdhury & Mohiuddin 2011; Khan, 2015a, 2015b). Among them, Seventy-six species from seven families have been reported from the north east region of Bangladesh (Khan 2015b). On the other hand, Ninety species have been reported from the south east region (Chowdhury & Mohiuddin 2011). However, the checklist of the eastern region is not comprehensive and many prospective habitats are yet to be explored.

The eastern region of Bangladesh is situated in the Indo-Burma Biodiversity Hotspot region and rich in diverse floral and faunal community. This region has a few semi evergreen forests and wildlife sanctuaries enriched with numerous streams and waterfalls. In addition to that, there are many marshes and lakes which provide ambient habitat for Odonates. However, despite being suitable habitat, till date scanty of studies have been carried out to annotate the Odonata fauna of the eastern region. Moreover, the previous research initiatives left many potential habitat to survey. The current study is a comprehensive approach for the documentation of the Odonata diversity of the eastern region of Bangladesh.

## 2.0 Materials and Methods

### 2.1 Study area

The odonates were surveyed the entire Sylhet division and the five districts e.g. Bandarban, Cox’s Bazar, Chittagong, Khagrachari and Rangamati of Chittagong division (Figure 1). In the north east region which is administratively under Sylhet division, Odonates were surveyed notably in Khadimnagar National Park, Tilagar Eco Park, Shahjalal University of Science and Technology campus, Satchari National Park, Lawachara National Park, Madhobpur Lake etc. On the other hand, in the south east region which is administratively under Chittagong division, Odonates were surveyed in the Chittagong University campus, Kaptai National Park, Bariadhala National Park, many streams and waterfalls associated areas of Chittagong, Khagrachari, and Bandarban district.

**Figure 1.**
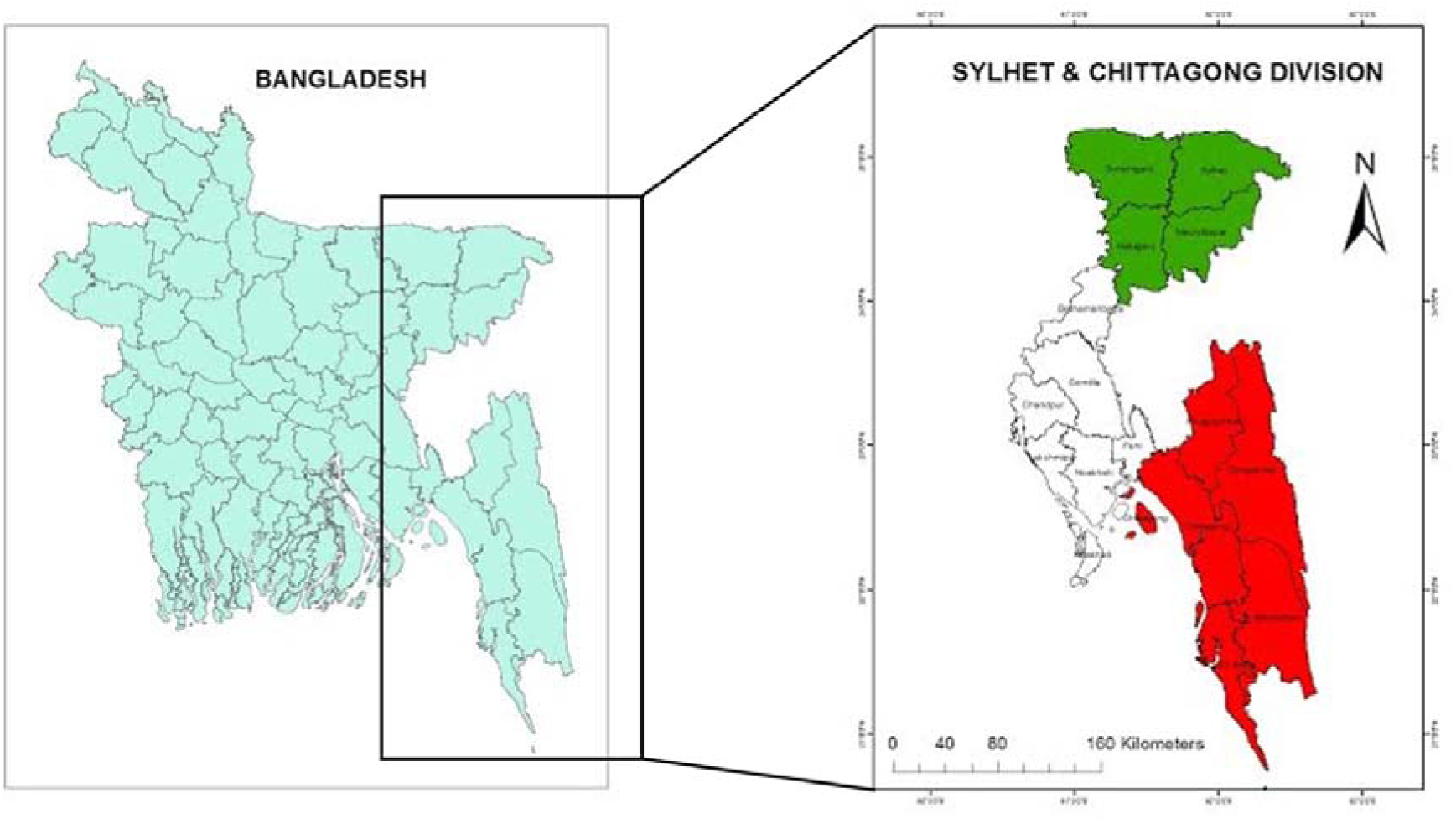
A reference map of the eastern region of Bangladesh. The north east region is administratively under Sylhet division and the south east is under Chittagong division. The red color and green color represents the area covered during the study period.

### 2.2 Specimen collection and identification

During the survey, the potential habitats like the marshes, ponds, streams, streams associated forest patches, temporary water sheds created during the monsoon were scanned thoroughly from 9.00 am to 4.00 pm. At field, the species were photographed for various identification keys using a Canon 600 DSLR camera (Canon Inc., Tokyo, Japan) fitted with a 55-250 mm telephoto zoom lens. The specimens were captured using an insect sweeping net and brought into the Department of Biochemistry and Molecular Biology, Shahjalal University of Science and Technology, Sylhet for further identification. In the laboratory, the specimens were examined under the microscope and identified based on the available identification keys provided by Fraser (1933, 1934, 1936) and Asahina (1993). The specimens were deposited in the Department of Biochemistry and Molecular Biology, Shahjalal University of Science and Technology, Sylhet, Bangladesh. The Odonates were classified according to Dijkstra et al. (2013).

### 3.0 Results

A total of 76 species from nine families belonging to 45 genera have been recorded from the eastern region of Bangladesh (Table 1; Figure 2). Among the documented Odonates, 46.05% (35 species) of 18 genera belong to Zygoptera sub-order while the rest 53.95% (41 species) of 27 genera belong to Anisoptera sub-order (Table 1). Libellulidae was the predominant Anisoptera family with 35 species from 22 genera (Table 1; Figure 2). On the other hand, Coenagrionidae was the best represented Zygoptera family with 19 species from 6 genera (Table 1; Figure 2). Four species (*Ceriagrion rubiae*, *Calicnemia imitans*, *Prodasineura autumnalis* and *Megalogomphus smithii)* have been recorded for the first time from Bangladesh.

**Figure 2.**
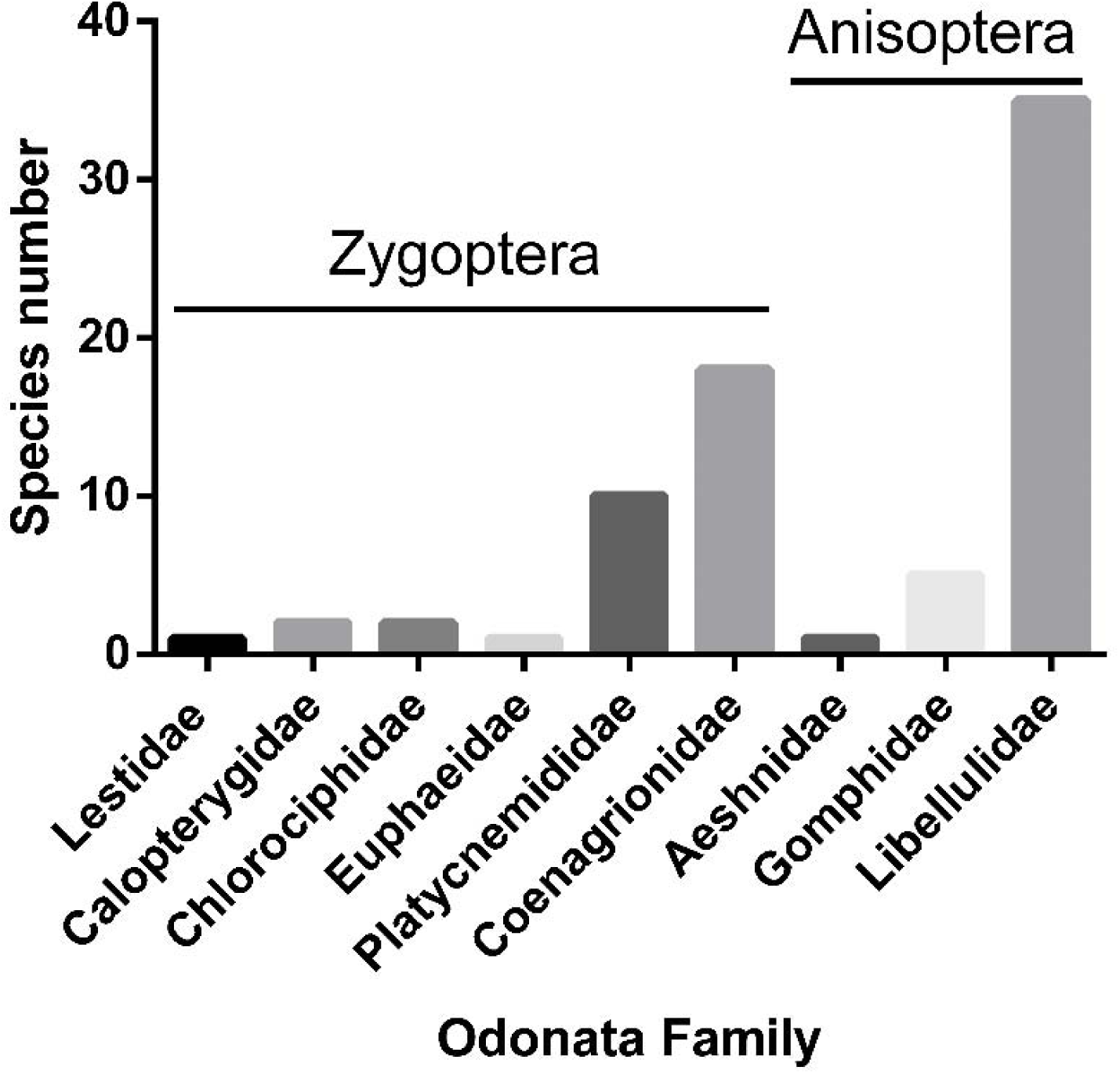
Number of Odonata species and their corresponding families recorded from the eastern region of Bangladesh during present study.

**Table 1:**
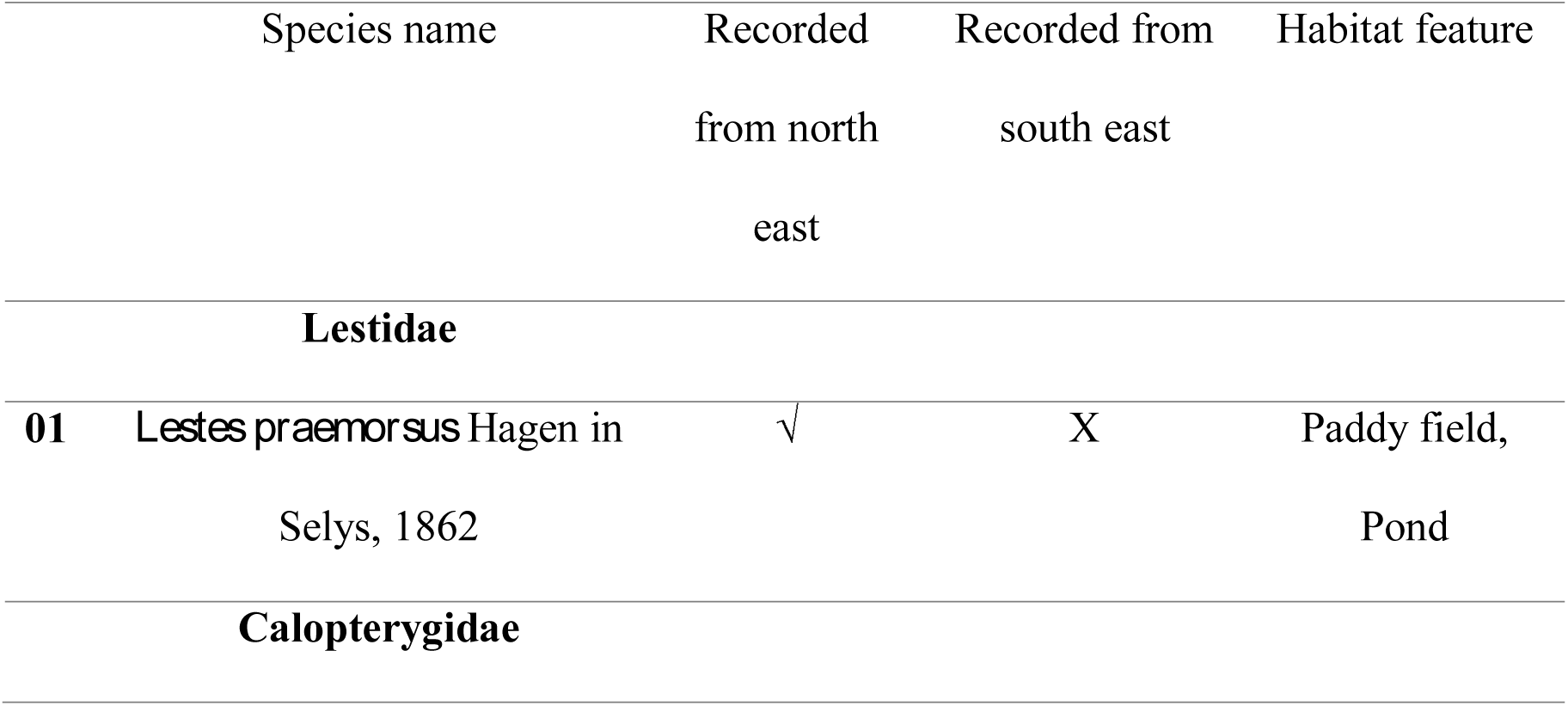
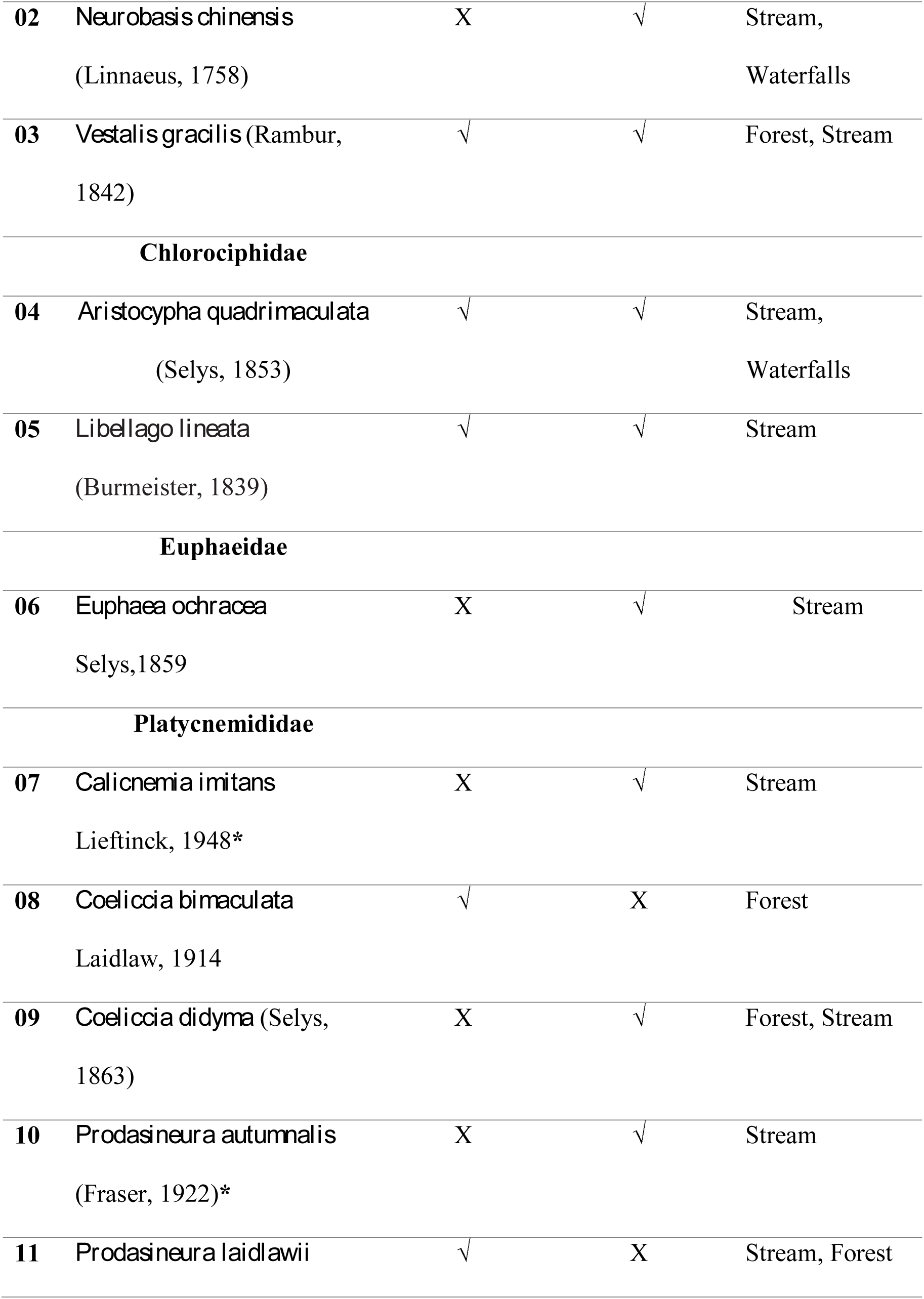
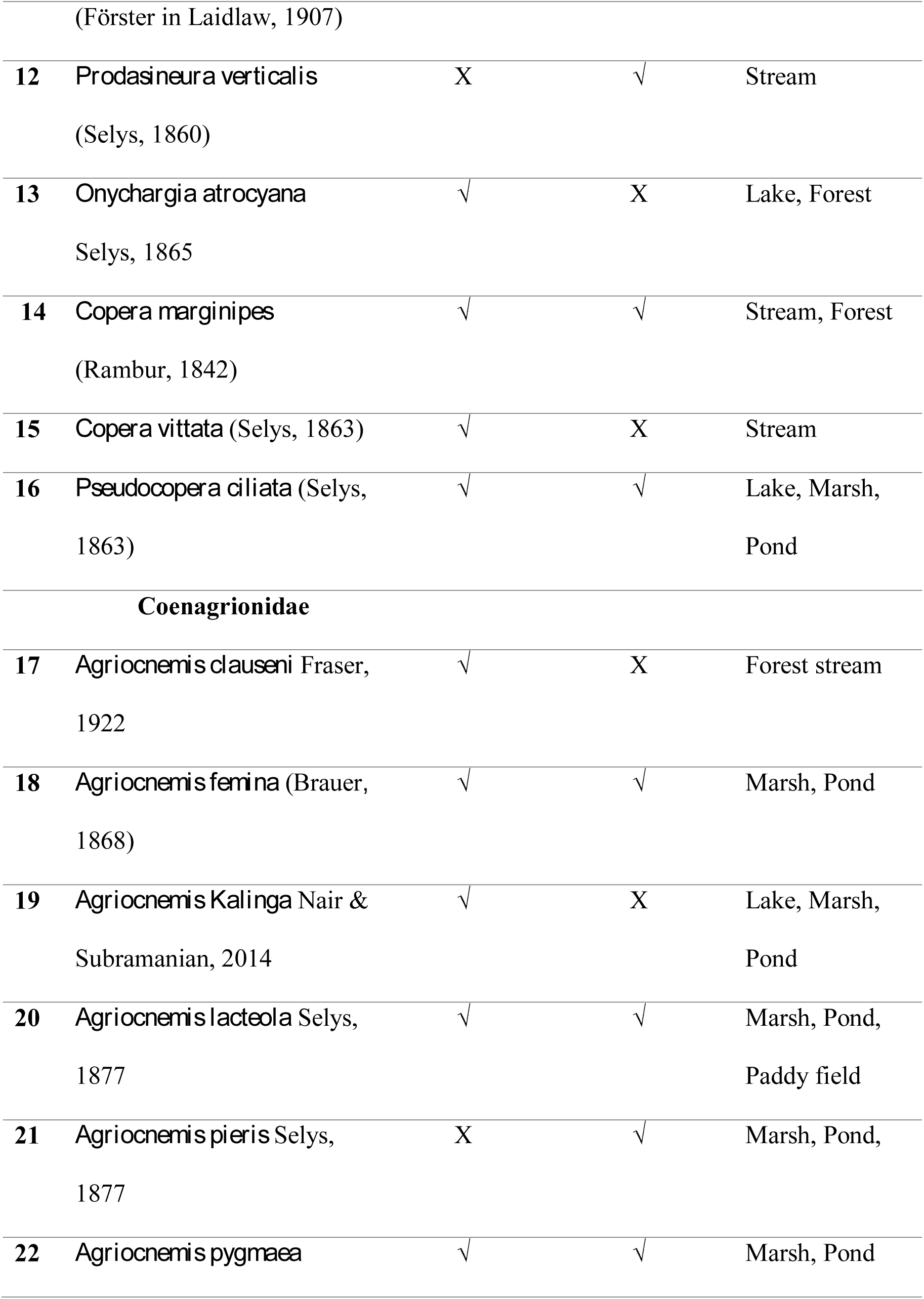
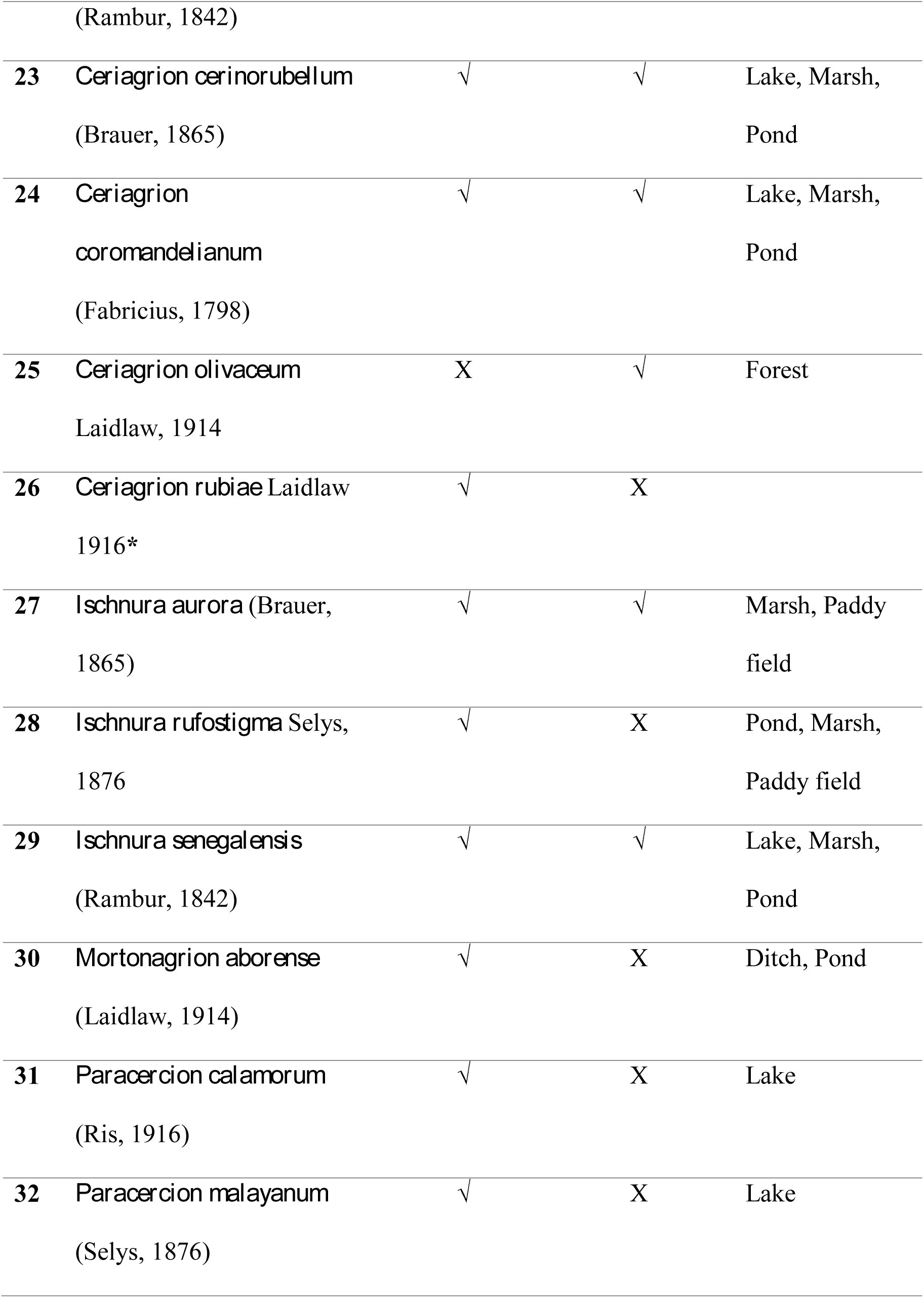
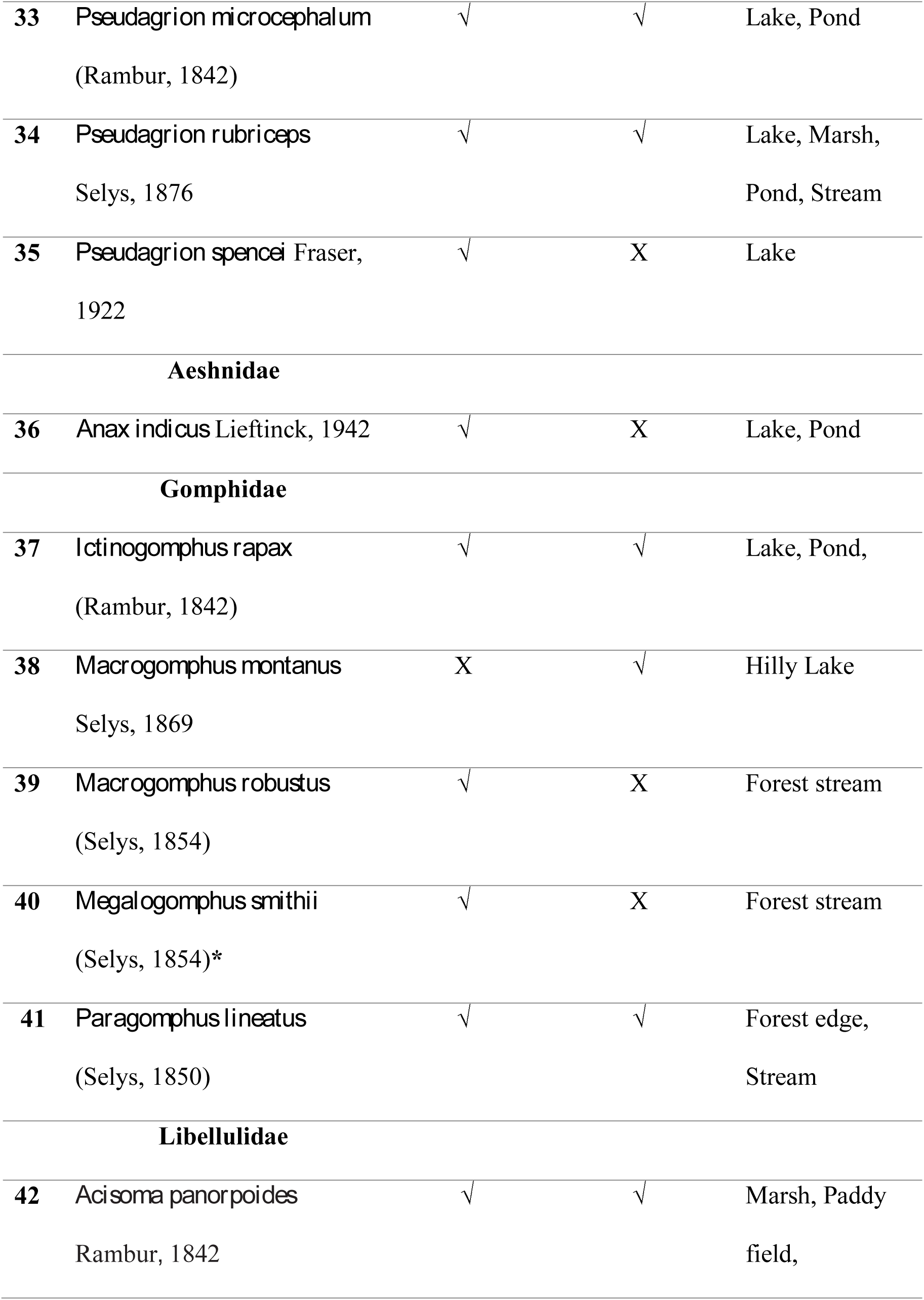
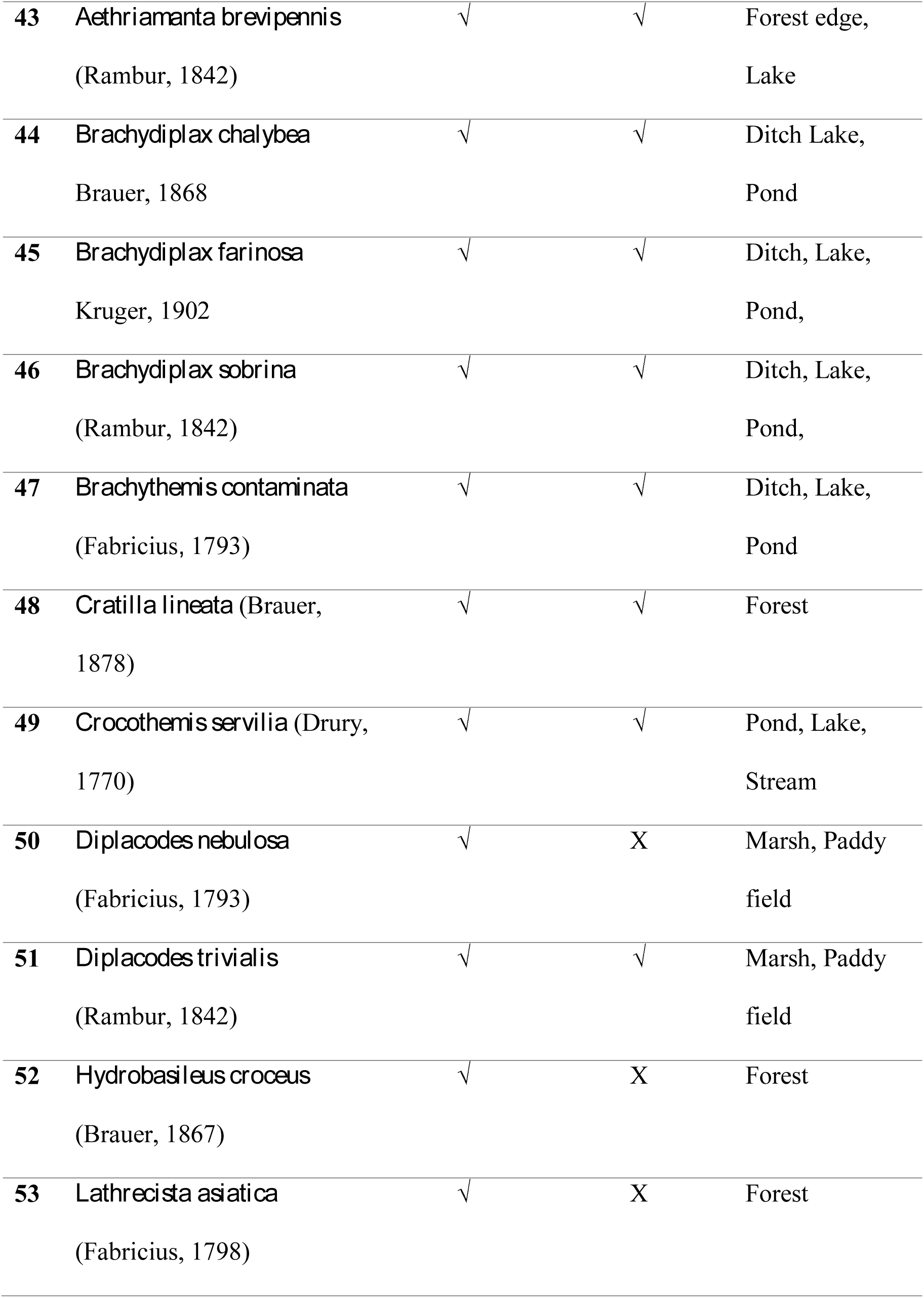
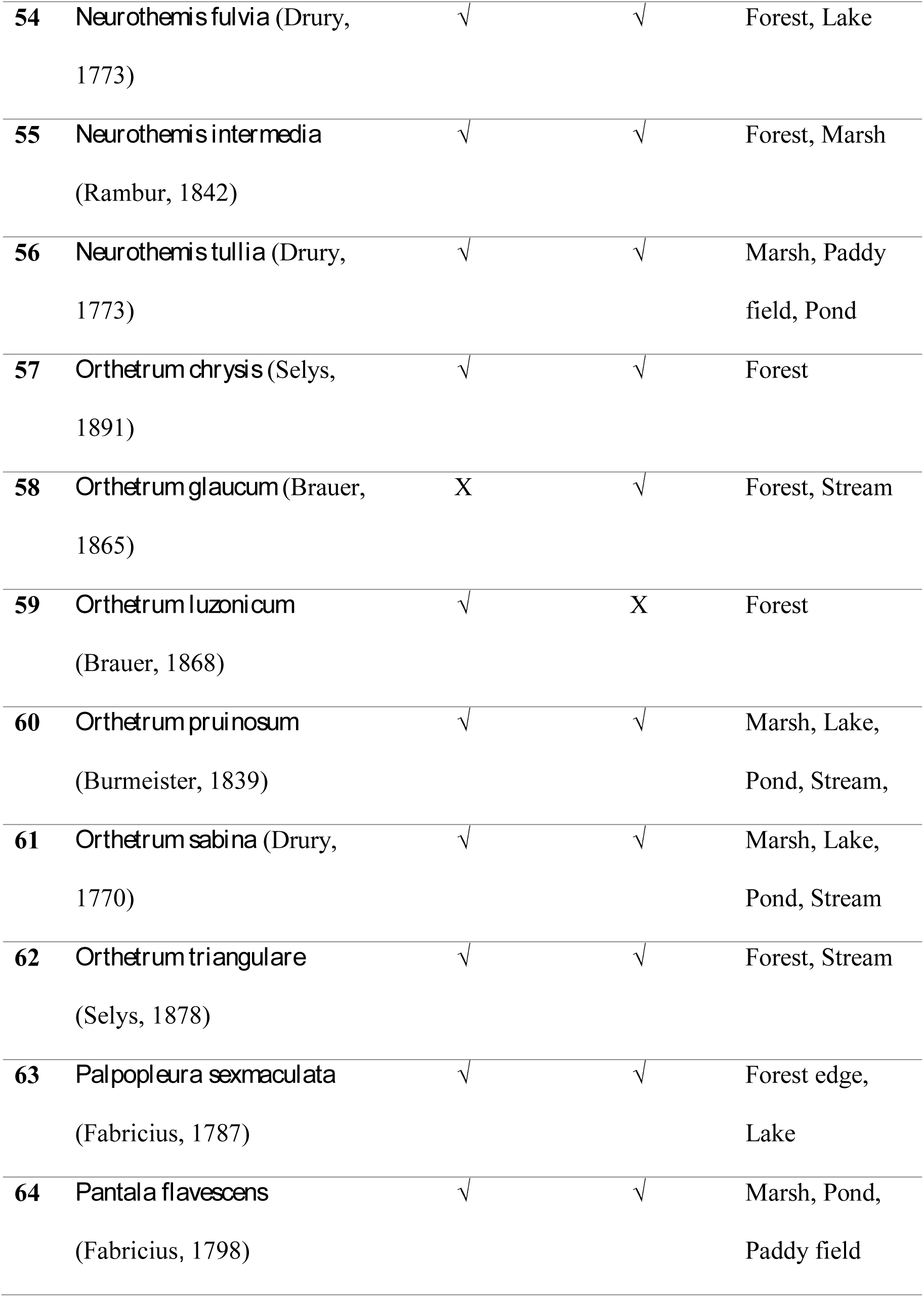
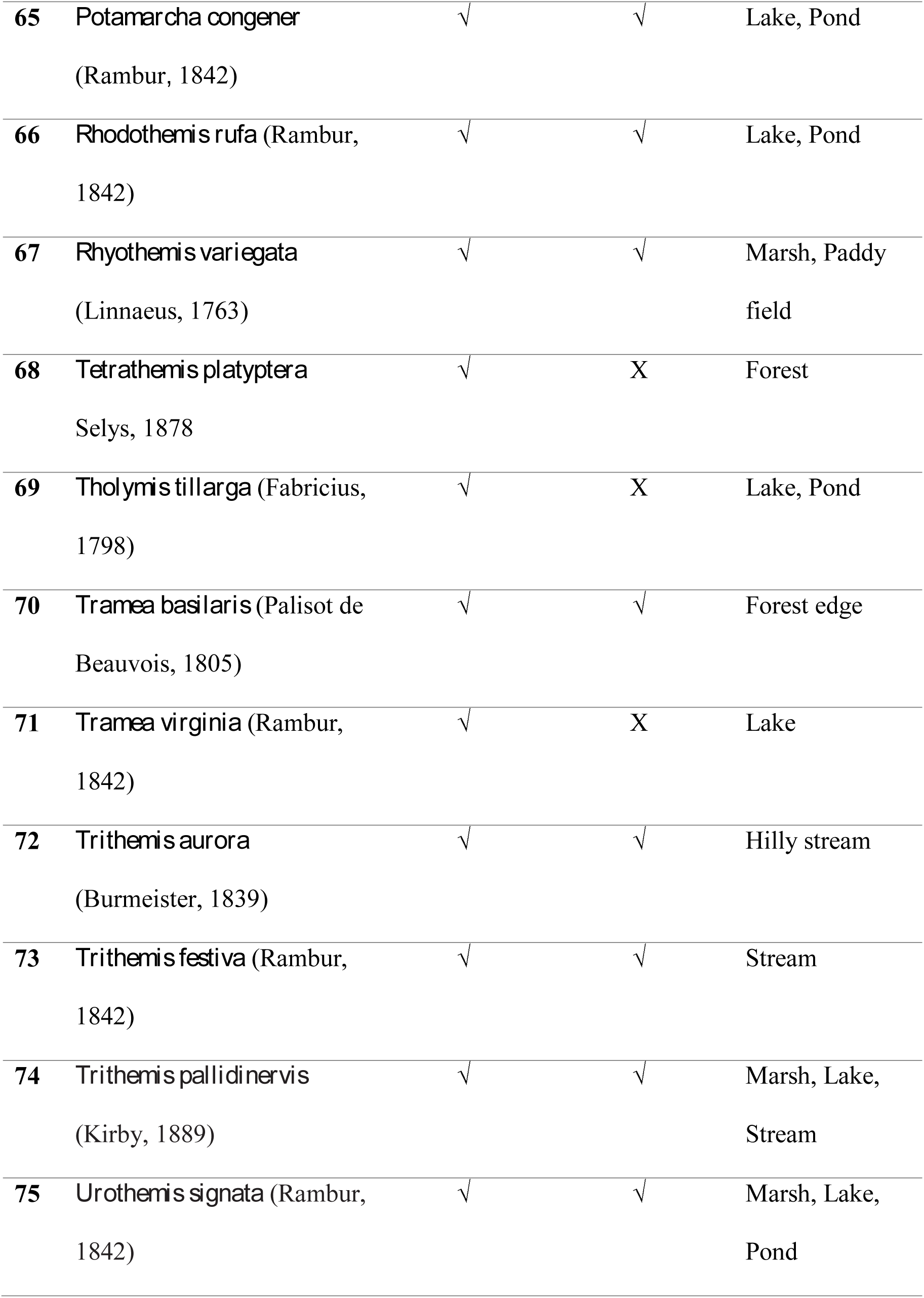
A list of the Anisoptera and Zygoptera species recorded in current study. The species newly discovered from Bangladesh are shown in asterisks (*). The species present in a particular area shown by tick sign (√) and the absent species are shown by cross mark (X).

A total of 66 species belonging to eight families have been recorded from the north east region. On the other hand, 52 species belonging to seven families have been documented from the south east region. Among the 76 recorded species, 41 species are found in the north east as well as in the south east region. Whereas, 24 species were recorded from the north east region only and 11 species were recorded from the south east part only. Coenagrionidae and Libelluidae were the best represented Zygopteran and Anisopteran family with 16 and 33 species respectively in the north east region. Similarly, in south east region Coenagrionidae and Libelluidae were the best represented Zygopteran and Anisopteran family with 11 and 27 species respectively.

### Newly recorded Odonates for Bangladesh

#### 1. *Ceriagrion rubiae* Laidlaw, 1916 (Figure 3A)

I have sighted *Ceriagrion rubiae* from the Shahjalal University of Science and Technology campus, Sylhet (24°92′112″N, 91°83′31″E) in June 2015. In May 2016, I have recorded this species again from the same location. The abdominal length and hind wing length of the male is 26-18 mm and 17-19 mm respectively. The male can be distinguished easily from the other Ceriagrion species by their bright, unmarked thorax and abdomen. Currently, this species is known from the geographical boundary of Bangaladesh and India only.

#### 2. *Calicnemia imitans* Lieftinck, 1948 (Figure 3B and 3C)

*Calicnemia imitans* is one of the most abundant species in the southeastern hilly streams. They prefer stream associated shaded bushes for perching. This is third recorded species of this genus from Bangladesh after *C. eximia* and *C. pulverulans*. I have recorded this species from the Alutila Cave, Khagrachari, Chittagong (23°05′18″N, 91°57′24″E) in June 2015. The length of the male abdomen is 29-31 mm and hind wing length is 20-22mm. This species can be distinguished from the other species by its body coloration and anal appendages. The ground color of male is black, orange and red color is absent in the thorax, narrow straight blue antehumeral stripe present, inferior is two third of the superior, tip of the superior is wide apart. This species is known from Bangladesh (New record), India, Laos, Myanmar, Thailand, Viet Nam (Fraser 1933; Hamalainen & Pinratana 1999; Cuong & Hoa 2007)

#### 3. *Prodasineura autumnalis* (Fraser, 1922) (Figure 3D, 3E)

I have recorded this species from the Kaptai National Park, Rangamati, Chittagong (22°29′50″N, 92°11′05″E) in October 2014. I have re-sighted this species later in June 2015 from Richang waterfalls, Khagrachari, Chittagong (23°06′38″N, 91°93′10″E), and Debota Pond, Khagrachari, Chittagong, (23°0'56"N, 91°58'18"E). The length of the abdomen and hind wing of the male is 30-31 mm and 18-20 mm respectively. *Prodasineura autumnalis* is superficially similar to *Prodasineura verticalis* and *Prodasineura sita*, however, they can be distinguished by the unmarked black thorax and the white tipped inferior anal appendages (Figure 3F). The females are found close to males and can be distinguished by blue antehumeral stripe (Figure 3 G). The species was previously known from China, India, Indonesia, Laos, Malaysia, Myanmar, Nepal, Singapore, Thailand and Vietnam (Fraser 1933; Vick 1989; Hamalainen & Pinratana 1999; Wilson & Reels 2003; Orr 2005; Cuong & Hoa 2007; Wilson 2005; Tang et al. 2010). The present record extened its distribution to Bangladesh also.

**Figure 3.**
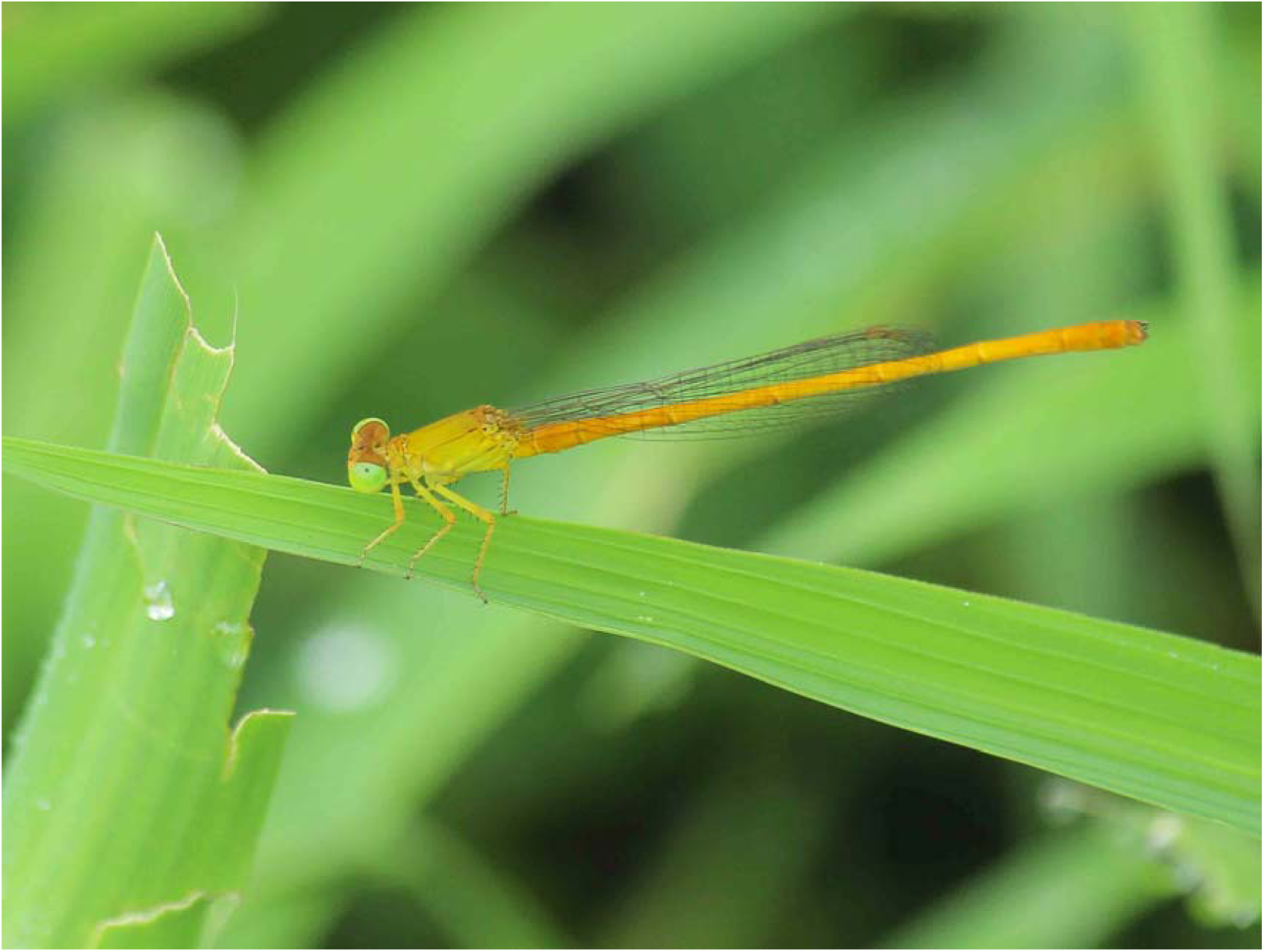
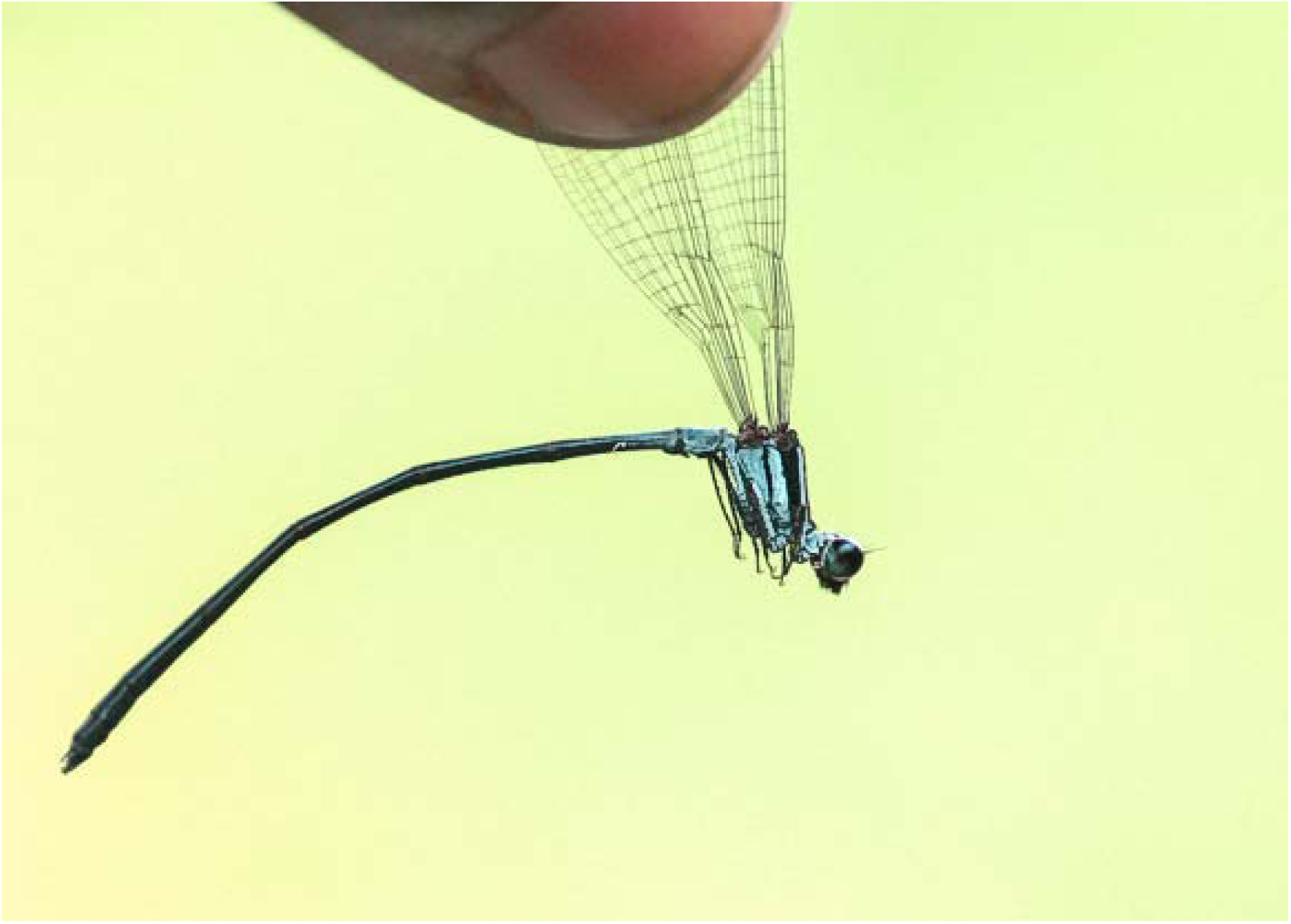
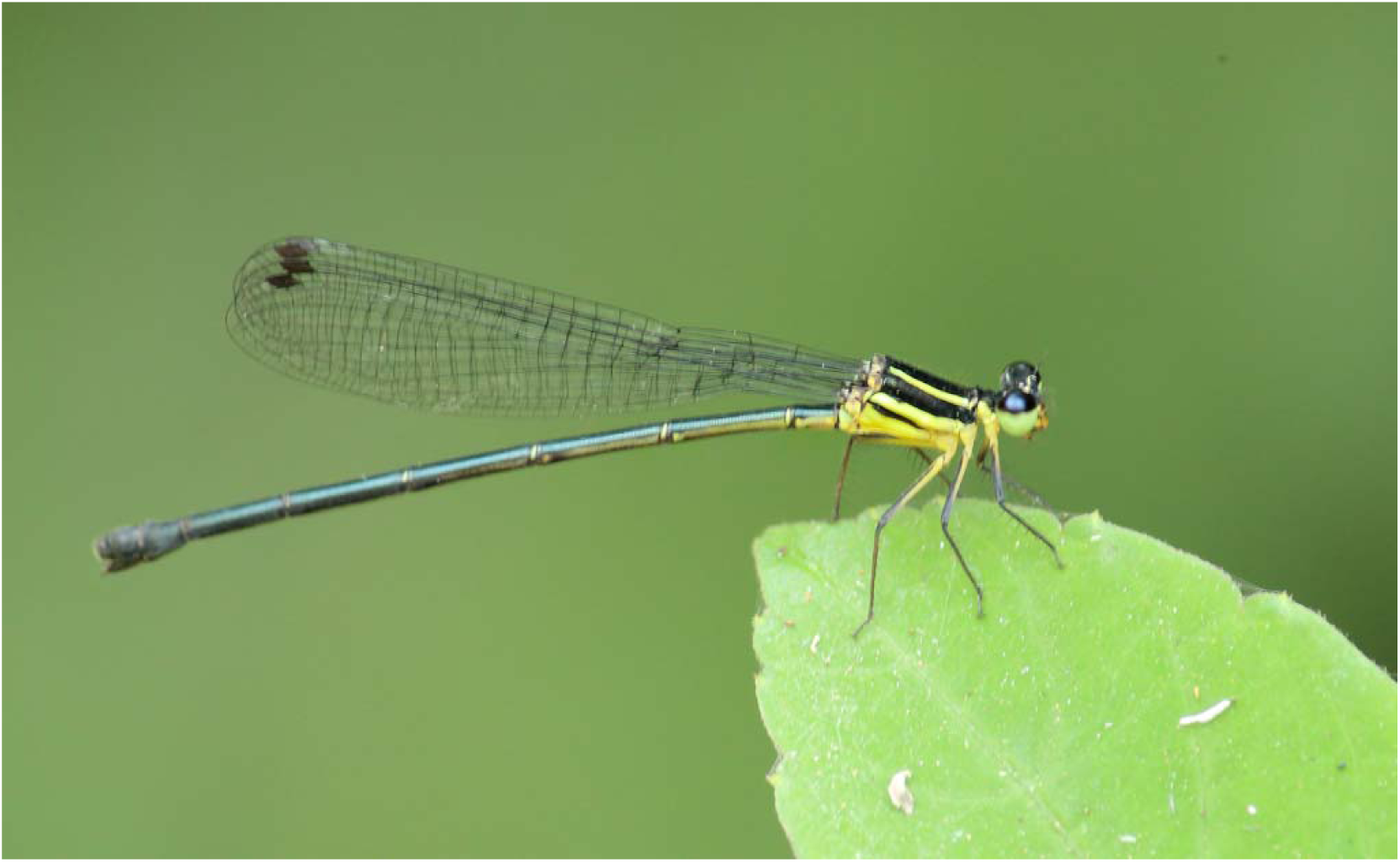
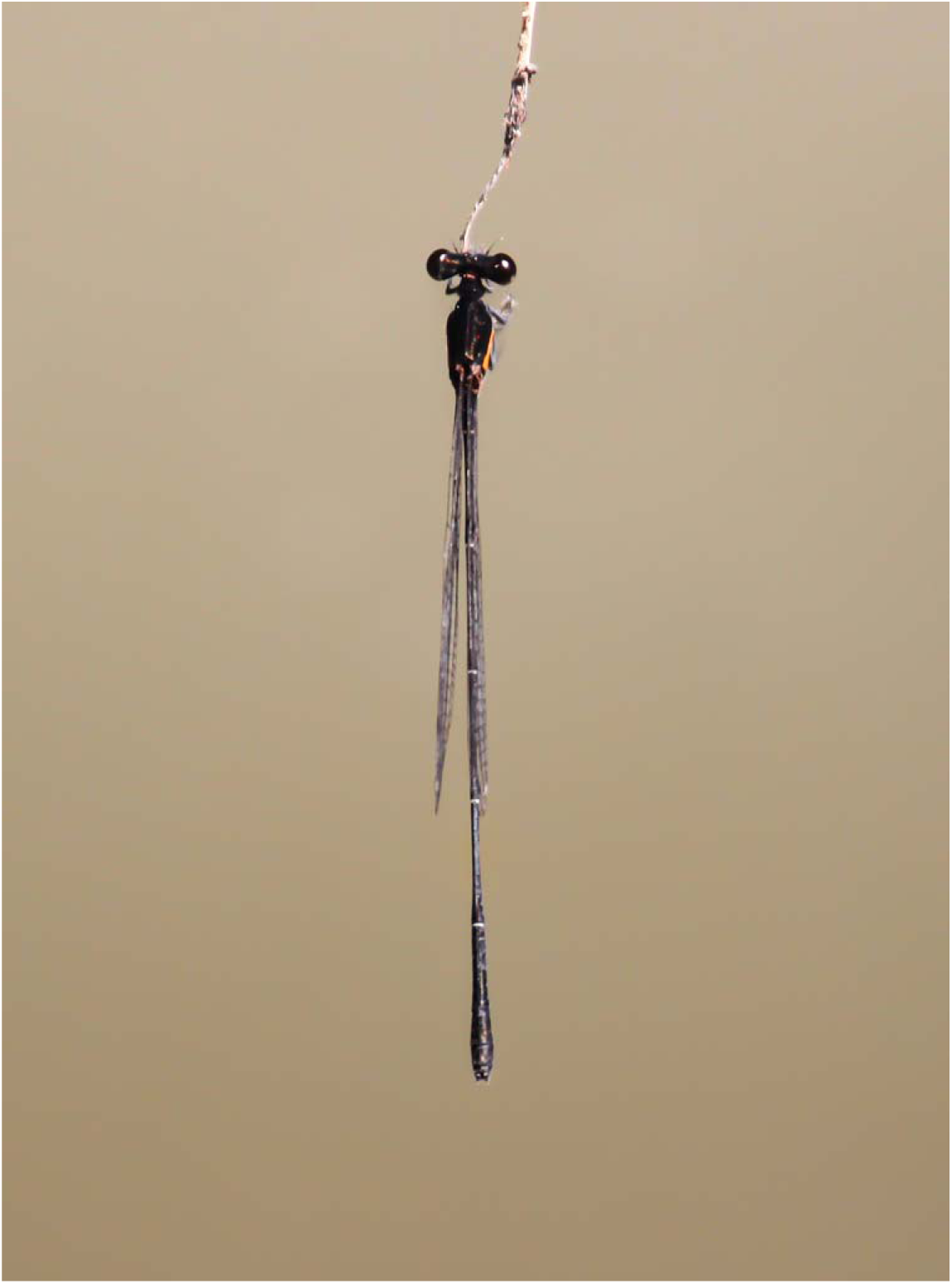
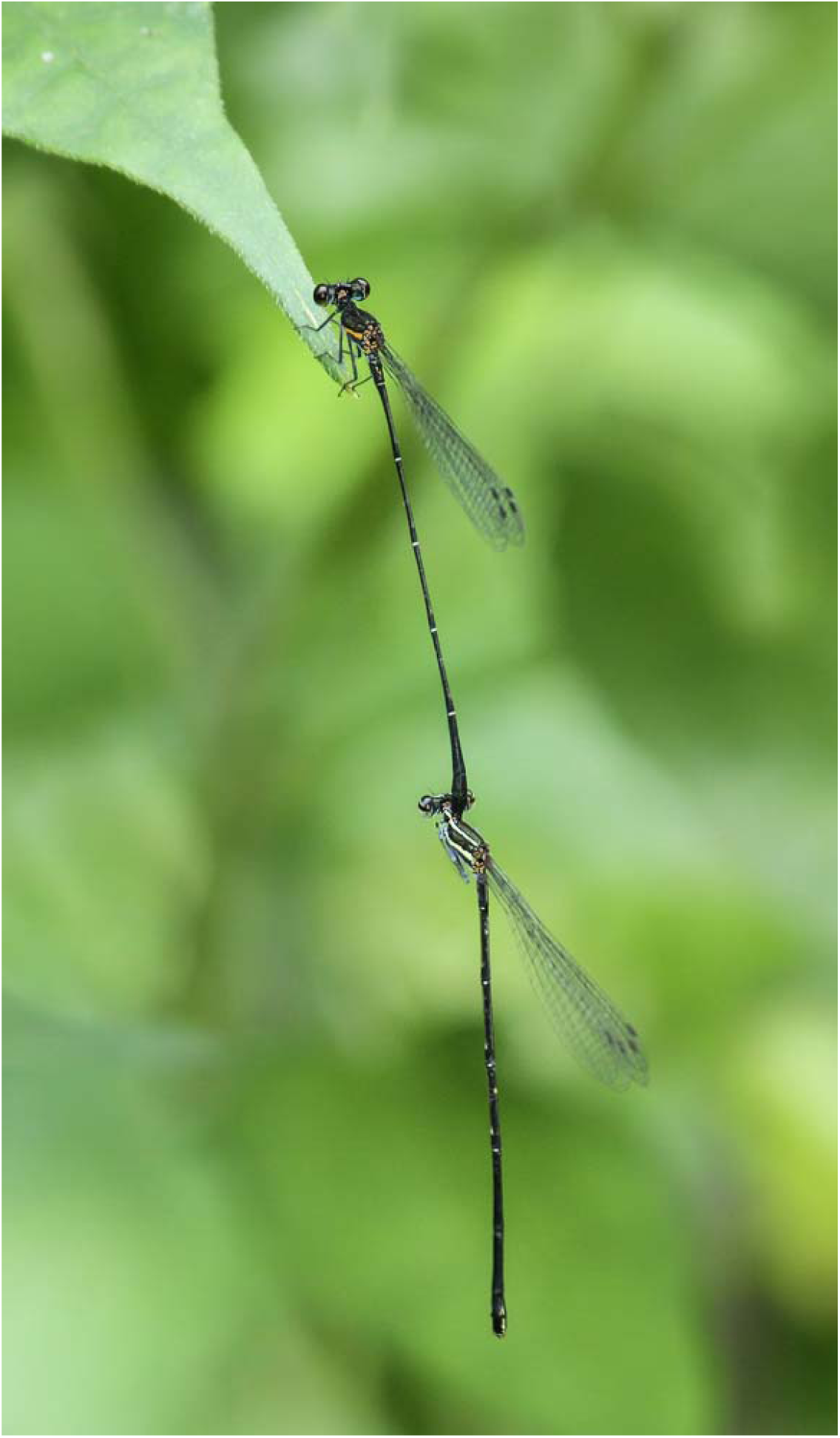
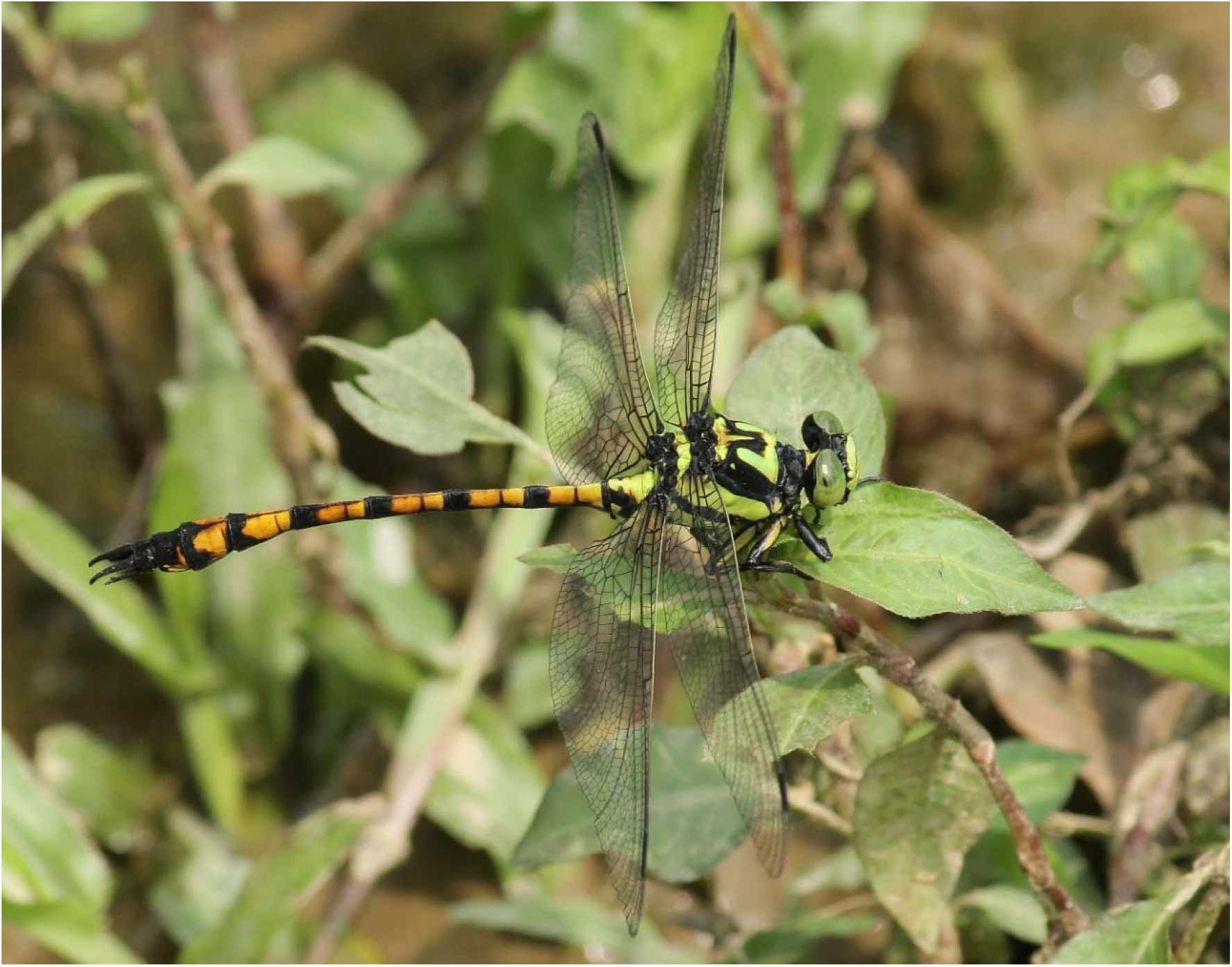
Photographs of the Zygoptera and Anisoptera species first time recorded from Bangladesh in present study. 3A. *Ceriagrion rubiae* Laidlaw, 1916 (male) 3B. *Calicnemia imitans* (male), 3C. *Calicnemia imitans* (female), 3D. *Prodasineura autumnalis* (male), and 3E. *Prodasineura autumnalis* (male and female in tandem position), 3F. *Megalogomphus smithii* (male)

#### 4 *Megalogomphus smithii* (Selys, 1854) (Figure 3F)

*Megalogomphus smithii* was previously known from Assam, India which is the adjacent to the north eastern region of Bangladesh. Considering to the similarity of habitats, this species was predicted to be present in Bangladesh also. I have recorded this species from Khadimnagar National Park Sylhet, Bangladesh (24°57′05″ N, 91°55′05″) in April 2015. The abdominal length of the male is 53-55 mm and the hind wing is 42-44 mm. This species has a prominent M shape marking in the thorax, and can easily be separated from the other member of the genus by the yellow marked black legs.

## 4.0 Discussion

In current study, Odonata fauna of the eastern region of Bangladesh have been documented. A total of 76 species from 40 genera have been recorded. Among them, four species and one genus have been recorded for the first time from Bangladesh. With the addition of this four species the current checklist of the Odonata fauna of Bangladesh is raised to 109 species. The high rate of the new record is a good indication that the Odonata fauna Bangladesh is poorly understood and demands more studies. Moreover, considering the habitat and Odonata fauna known from adjacent states of India e.g. Assam, Meghalaya, West Bengal and Myanmar, it can be predicted that more Odonata species should be present in Bangladesh.

Regional checklist is a good indicator of the diversity of particular faunal community, their distribution range, and population fragmentation. Hence, updating regional checklist on a regular basis is a good practice to understand the conservational status of a species. In current study, three species (*Agriocnemis clauseni, Pseudagrion spencei* and *Tramea Virginia*) are newly added to the odonata fauna of the north east region. In addition to that, the current study has extended distribution range of a few previously recorded species. The distribution range extension and new habitat allocation is particularly important to assesses the national status of those species and also for the IUCN Red Listed globally data deficient, endemic and endangered species. In current study, distribution range has been extended for two globally data deficient species. Among them, *Macrogomphus robustus* have been previously recorded from Lawachara National Park, Maulavibazar (Chowdhury & Mohiuddin 2011). The present record extends its distribution further north to the Khadimnagar National Park, Sylhet. The other data deficient species, *Megalogomphus smithii* have been previously known from China, India, and Indonesia. The present study reported this species for the first time within the geographical area Bangladesh. The individual number of this two data deficient species recorded from the current study is very low and thus long-term studies are essential to assess their population trends and distribution range.

In conclusion, the diverse Odonata fauna of the eastern region and newly recorded species of the area indicates it may accommodate hitherto unknown species. Moreover, the current study suggests that more long-term surveys are required to annotate the Odonata fauna of Bangladesh, to estimate their current status and to determine conservation needs.

## Acknowledgements

I am very thankful to two anonymous reviewer for their comments on the initial version of the manuscript. I am also thankful to Payal Barua, Tabrakullah Ranu, Nur Ahad Shah, Shuvo Sutradhar, Rumana yesmin Trisha and Rakhal Chandra Das for their support during the field study. I would also like to acknowledge Noppadon Makbun and Rupa Saha for their support during in taxonomy and in manuscript preparation respectively.

